# Elucidating genomic patterns and recombination events in plant cybrid mitochondria

**DOI:** 10.1101/506816

**Authors:** Laura E. Garcia, Mikhajlo K. Zubko, Elena I. Zubko, M. Virginia Sanchez-Puerta

**Affiliations:** IBAM, Universidad Nacional de Cuyo, CONICET, Facultad de Ciencias Agrarias, Almirante Brown 500, M5528AHB, Chacras de Coria, Argentina & Facultad de Ciencias Exactas y Naturales, Universidad Nacional de Cuyo, 5500, Mendoza, Argentina; Faculty of Science and Engineering, Manchester Metropolitan University, Manchester, M1 5GD, UK

**Keywords:** mitochondrial DNA, cybrid, DNA recombination, BIR, protoplast fusion

## Abstract

The maintenance of a dynamic equilibrium between the mitochondrial and nuclear genomes requires continuous communication and a high level of compatibility between them, so that alterations in one genetic compartment need adjustments in the other. The co-evolution of nuclear and mitochondrial genomes has been poorly studied, even though the consequences and effects of this interaction are highly relevant for human health, as well as for crop improvement programs and for genetic engineering. The mitochondria of plants represent an excellent system to understand the mechanisms of genomic rearrangements, chimeric gene formation, incompatibility between nucleus and cytoplasm, and horizontal gene transfer. We carried out detailed analyses of the mitochondrial genome (mtDNA) of a repeated cybrid between the solanaceae *Nicotiana tabacum* and *Hyoscyamus niger*. The mtDNA of the cybrid was intermediate between the size of parental mtDNAs and the sum of them. Noticeably, most of the homologous sequences inherited from both parents were lost. In contrast, the majority of the sequences exclusive of a single parent were maintained. The mitochondrial gene content included a majority of *N. tabacum* derived genes, but also chimeric, two-parent derived, and *H. niger-derived* genes in a tobacco nuclear background. Any of these alterations in the gene content could be the cause of CMS in the cybrid. The parental mtDNAs interacted through 28 homologous recombination events and a single case of illegitimate recombination. Three main homologous recombination mechanisms were recognized in the cybrid mitochondria. Break induced replication (BIR) pathway was the most frequent. We propose that BIR could be one of the mechanisms responsible for the loss of the majority of the repeated regions derived from *H. niger*.

## Introduction

Mitochondria are semi-autonomous cellular compartments, in which numerous essential metabolic pathways take place (Logan, 2007; Taanman, 1999). This organelle contains its own genome and molecular machinery for gene expression. However, the maintenance of the mitochondrial genome (mtDNA) and the transcription of mitochondrial-encoded genes are controlled by the nucleus. Therefore, mtDNA replication, structural organization, and gene expression depend on the coordinated interaction between the mitochondrial and the nuclear genomes (Chase, 2007). The effects of the co-evolution of these genomes are relevant for the human health (Chou and Leu, 2015), for breeding programs (Orczyk et al., 2003), and in genetic engineering (Greiner and Bock, 2013). Angiosperm mtDNAs have particular characteristics that distinguish them from those in fungi and animals: very low substitution rates, highly scrambled gene order, relatively large and variable genome sizes (between 222 kb to 11.3 Mb), and long intergenic regions that encompass up to 98% of the genome (Kubo and Mikami, 2006; Skippington et al., 2015; Sloan et al., 2012). A part of these intergenic regions could be attributed to repeats and to sequences acquired from the nuclear and chloroplast genomes by intracellular gene transfer or from other plant mitochondria via horizontal gene transfer (Rice et al., 2013; Sanchez-Puerta et al., 2008; Xi et al., 2013), while the major part remains to be characterized (Gualberto and Newton, 2017). Frequent homologous recombination events across repeats make plant mtDNA prone to rearrangements, evolving rapidly in structure (Arrieta-Montiel, and Mackenzie, 2011; Palmer and Herbon, 1988). The mixing of mtDNA regions by homologous recombination can modify gene organization and create chimeric genes that play a role in evolution (Gualberto and Newton, 2017). In particular, the cytoplasmic male sterility (CMS) is an economically important trait that can result from the formation of chimeric mitochondrial genes (Chase, 2007; Touzet and Meyer, 2014). The generation of somatic hybrids via protoplast fusions has been used for years to introduce CMS to crop species, as well as to accelerate and improve various aspects of plant breeding (Gleba and Sytnik, 1984; Orczyk et al., 2003).

Somatic hybridization between different plant species represents a powerful method to generate novel traits or to transfer organelle-encoded features to cultivars. Cybrids (cytoplasmic hybrids) are somatic hybrids that contain the nuclear genome of one parent while the cytoplasmic genomes could be inherited from the other parent or from both parents. The production of cybrids, in particular, is an important tool for crop improvement and to study nuclear-cytoplasmic interactions and evolutionary dynamics (Greiner and Bock, 2013; Woodson and Chory, 2008). Somatic hybrid production has been widely used in Solanaceae breeding programs (Austin et al., 1988; Brown et al., 1996; Iovene et al., 2007; McGrath et al., 2002; Orczyk et al., 2003). Interspecific and intergeneric somatic and cytoplasmic hybrids have been produced between *Nicotiana* spp. and related species to introduce relevant traits, such as CMS (Carlson et al., 1972; Fitter et al., 2005; Ilcheva et al., 2000; Sun et al., 2005).

It is well known that plant somatic hybridization often results in recombinant mitochondrial genomes (Babiychuk et al., 1995; Belliard et al., 1979; Gleba and Sytnik, 1984; Vedel et al., 1986; Zubko et al., 1996) Characterization of these chimeric mitogenomes has been quite limited, including the identification of a small number of parental-specific markers by Southern blot hybridizations (Aviv et al., 1984; Belliard et al., 1979; Kofer et al., 1991; Morgan and Maliga, 1987; Nagy et al., 1983; Rothenberg and Hanson, 1988; Scotti et al., 2004) and the complete sequence of two cybrid mtDNAs (Arimura et al., 2018; Sanchez-Puerta et al., 2008). Genomic comparisons between parental and cybrid cytoplasmic genomes showed that the cybrid mitochondrial genomes were larger than each parental mtDNA and remarkably chimeric. Despite their increased size, the cybrid mtDNAs contained single alleles for most of the mitochondrial genes. Retaining a single form of each gene could be a selective mechanism to minimize intracellular incompatibilities or a neutral process that preferentially eliminates duplicated regions (Sanchez-Puerta et al., 2015). Further analysis of hybrid mitochondria would help to elucidate the nature and evolutionary dynamics of the molecular processes in plant mitochondria, such as genomic recombination, nuclear-cytoplasmic interactions, and the incorporation of foreign sequences through horizontal gene transfer.

In this study, we aim to gain insight into the mitochondrial genome dynamics of cybrid plants through sequencing and analysis of the mtDNA of a repeated cybrid between *Nicotiana tabacum* and *Hyoscyamus niger*. This cybrid combination was chosen because both parental mtDNAs were sequenced (Gurdon et al., 2016; Sanchez-Puerta et al., 2015; Sugiyama et al., 2005) and a previous study on a cybrid produced from the same parent combination was reported (Sanchez-Puerta et al., 2015), allowing a direct comparison of the evolutionary path taken by this independent cybrid. We wish to address the following questions: Were the homologous sequences from each parent mtDNA lost from the cybrid mitochondria as observed in the *N. tabacum (+H. niger)* cybrid previously analyzed? Did the cybrid mtDNA maintain the sequences that are exclusive of each parental mitochondria? Has the initial co-existence of the two parental alleles in the cybrid mtDNA been maintained? Is there a strong preference for tobacco-derived intergenic and/or genic sequences given that the nuclear genome was inherited from tobacco?

## Materials and Methods

### Genome Sequencing, assembly and validation

Total DNA was extracted from leaves of an individual plant (Mv-1-1g) of the repeated cybrid line Drhn-3 between *Nicotiana tabacum* and *Hyoscyamus niger* (Figure 1) following the Dellaporta method (Dellaporta et al., 1983). It was used to construct a paired end library with insert size of ~800 bp that was sequenced at the Beijing Genomics Institute using the Illumina Hiseq 2500 sequencing system, generating 70 million clean paired-end reads of 125 bp. For the reconstruction of the mtDNA we carried out the following methodology. First, we performed *de novo* assembly using Velvet v. 1.2.03 (Daniel Zerbino, European Bioinformatics Institute Cambridge, UK) on the Mason large-memory computer cluster at Indiana University-Bloomington (IN, USA). We ran eight independent Velvet *de novo* assemblies without scaffolding and using different hash lengths each time (k-mer lengths: 41, 73, 81, 97, 99, 103, 107, 111). The best assembly (k-mer 103) resulted in 387 contigs larger than 1 kb with N50 contig length of 5,249 bp and 35 contigs in the N50. Using BLAST we selected 66 contigs that matched the mitochondrial genomes of the parents (*H. niger* and *N. tabacum*) and had query coverage (qcov) greater than 50%. These putative-mitochondrial contigs were used as trusty contigs in the assembler SPAdes (Bankevich et al., 2012), producing 51 contigs that were later extended using SSAKE (Warren et al., 2007). The extended contigs were joined using SSPACE (Boetzer et al., 2011) and Gapfiller (Boetzer and Pirovano, 2012). Finally the resulting contigs were manually assembled based on consistent paired-end reads using Consed v.26 (Gordon and Green, 2013). The quality of mitochondrial genome assembly was evaluated by mapping all Illumina reads until the read depth was as expected and the high-quality mismatches were very low or zero. The DNA library had a mean size estimated by Consed v.26 of 769 bp, SD 139. Repeats shorter than 750 bp and with >80% identity were identified with crossmatch in Consed v.26. To identify repeats larger than the insert size, we took into account the total read depth and paired-end read information using Consed v.26. To confirm the absence of complete mtDNA sequences from the parents (even at low stoichiometries) in the cybrid mitochondria, we mapped all Illumina reads to the *H. niger* or *N. tabacum* mitochondrial genome sequences using Consed v.26. The total and paired-end read depth across the parental mtDNAs was irregular, showing regions that lack reads and regions with inconsistent paired-end reads. Thus, we confirmed that the cybrid mitochondria did not contain the full-length genomes from its parents but it was composed of a subset of sequences from both parents. Chloroplast contigs were identified through BLAST and the chloroplast genome was assembled using Novoplasty (Dierckxsens et al., 2017).

**Figure 1.**
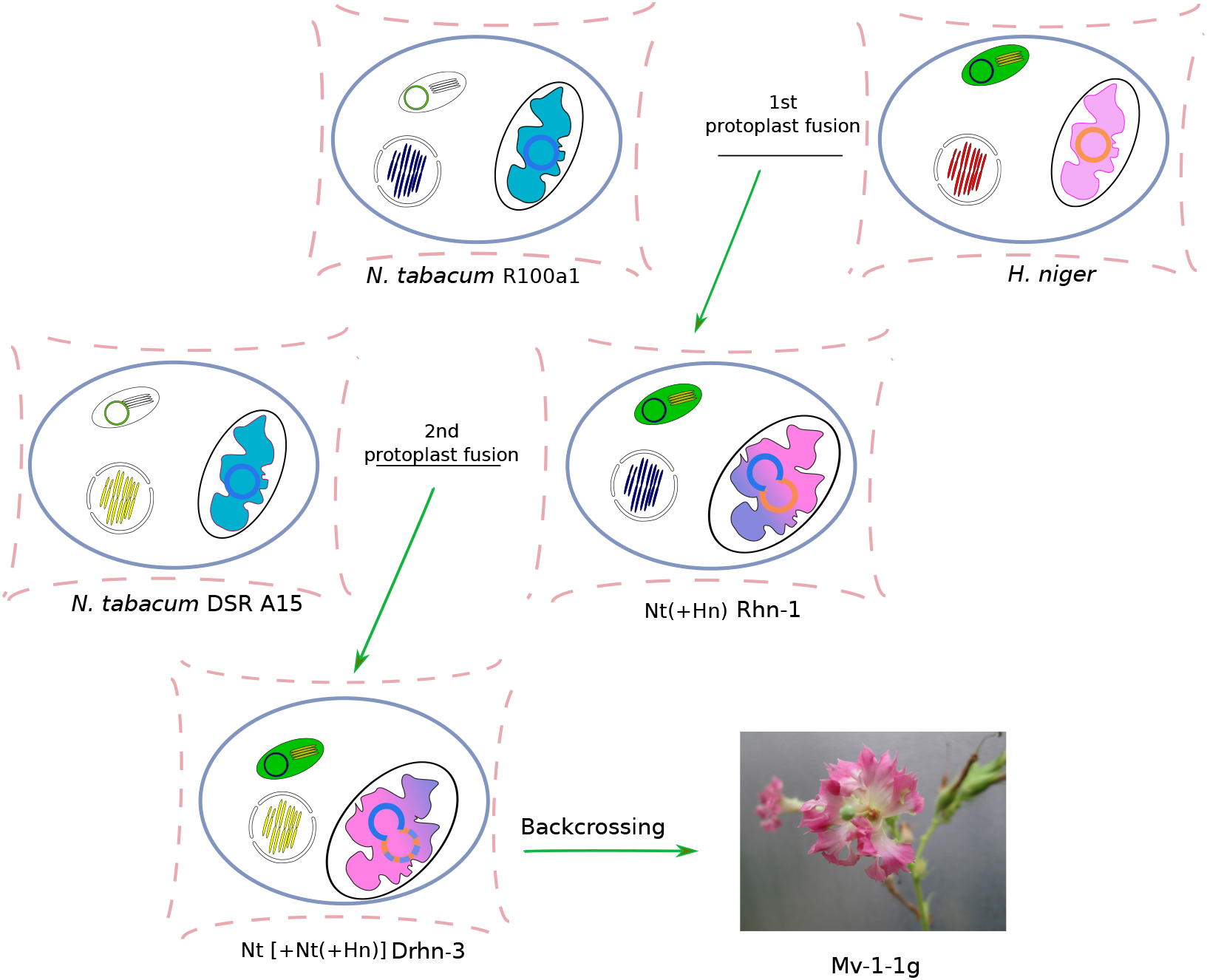
Schematic production of the repeated cybrid plant Mv-1-1g in a two-step protoplast fusion experiment. In the first step, protoplasts from an albino mutant line of *Nicotiana tabacum* and a wild-type line of *Hyoscyamus niger* were fused. Among green hybrid plants there was a cybrid line Rhn-1 manifesting a pistilloid form of CMS. Protoplasts of Rhn-1 were inactivated by gamma irradiation before the second protoplast fusion with tobacco plastome mutant DSR A15. Mv-1-1g is a backcross descendant of the line Drhn-3 with wild-type *N. tabacum*.

All raw sequence data are available from the NCBI Sequence Read Archive database as accession number xxxxxxx. The annotated cybrid mitochondrial genome was deposited in the GenBank data libraries under accession numbers MH168702- MH168706.

### Genome annotation

Mitochondrial contigs were annotated using the software Geneious v. 11.0.4, BLAST (Altschul et al., 1990), and tRNAscan (Lowe and Chan, 2016). Graphical genome maps were generated using OGDraw (Lohse et al., 2007). Unknown ORFs (>300 bp and with ATG as start codon) were identified using Geneious v. 11.0.4.

### Comparisons between genomes

BLASTn comparisons of the mitochondrial genomes of the cybrid and its parents (*N. tabacum:* KR780036, NC_006581 and *H. niger*: NC_026515) were visualized in Genome Workbench and Geneious v.11.0.4. Putative intergenomic recombination events in the cybrid mtDNA were inferred by visual inspection of BLAST results. Gene conversion events (without crossover) were inferred when three or more polymorphisms in a short tract (< 400 bp) were shared by the cybrid and one parent inside a region that was otherwise clearly derived from the other parent. Genes were aligned, and differences between *N. tabacum* and *H. niger* in protein-coding genes were scored as synonymous or nonsynonymous substitutions using Geneious v.11.0.4.

## Results

### Cybrid production

The Mv-1-1g repeated cybrid was constructed by a two-step protoplast fusion involving two species of the family Solanaceae (Figure 1). In the first experiment, protoplasts from a recipient line of *Nicotiana tabacum* L. 2n=4x=48 plastome albino mutant (R100a1) were fused with those from a donor line of wild type *Hyoscyamus niger* L., 2n =2x=34 (Zubko et al., 1996). This fusion event resulted in the cybrid line Rhn-1 that was phenotypically similar to tobacco with the prevalence of the tobacco nuclear genome after 4 years of vegetative propagation. The cybrid exhibited cytoplasmic male sterility (CMS) and developed flowers without corolla and stamens that, instead, had a complex of 3-4 flower-like structures with pronounced pistilloidy (Zubko et al., 1996). The cytoplasm of Rhn-1 was re-transferred into the nuclear background of a plastome chlorophyll deficient-tobacco mutant (DSR A15) in a second experiment of protoplast fusion (involving inactivation of Rhn-1 protoplasts with gamma-irradiation) and posterior *in vitro* culture (Figure 1). One of the resulting cybrid lines was named Drhn-3. The plants of this line rooted *in vitro*, were transferred to soil and placed in a greenhouse for repeated backcrosses with wild type tobacco. Drhn-3 plants exhibited CMS, wrinkled corollas, no stamens, and one pistil. Here, we studied a cybrid plant (named Mv-1-1g) that was obtained after two backcrosses of the cybrid line Drhn-3 with wild type tobacco. This plant manifested more elongated leaves (in comparison to wild-type tobacco), funnel-like corolla (in contrast to tobacco bell-like corolla), crimpy petals, and CMS (due to undeveloped stamens without pollen). The chloroplast genome of the cybrid Mv-1-1g was inherited from *H. niger* and 100% identical to the *H. niger* cpDNA.

### The cybrid Mv-1-1g Mitochondrial Genome

We sequenced and assembled the mitochondrial genome (mtDNA) of the Mv-1-1g repeated cybrid. The mtDNA assembled into 5 contigs (ranging between 9,520 and 359,619 bp) with a total length of 707,034 bp (Figure 2). The genome length was intermediate between the size of parental mtDNAs (~501 and ~431 kb) and the sum of them (~932 kb). The total read depth was fairly even across the genome (Suppl. Figure 1). All mitochondrial contigs could potentially recombine with each other through large and intermediate repeats. There were 18 large (>1 kb), 16 intermediate (250-1,000 bp), and 81 short (100-250 bp) repeats in the cybrid mtDNA. Also, reads at the end of each contig had their mate reads in a distant region of the same or other contig, indicating the existence of alternative configurations of the cybrid mitochondrial genome. The GC content of the genome was 44.6%, ranging between 44.2 and 44.9 *%* in each individual contig.

**Figure 2.**
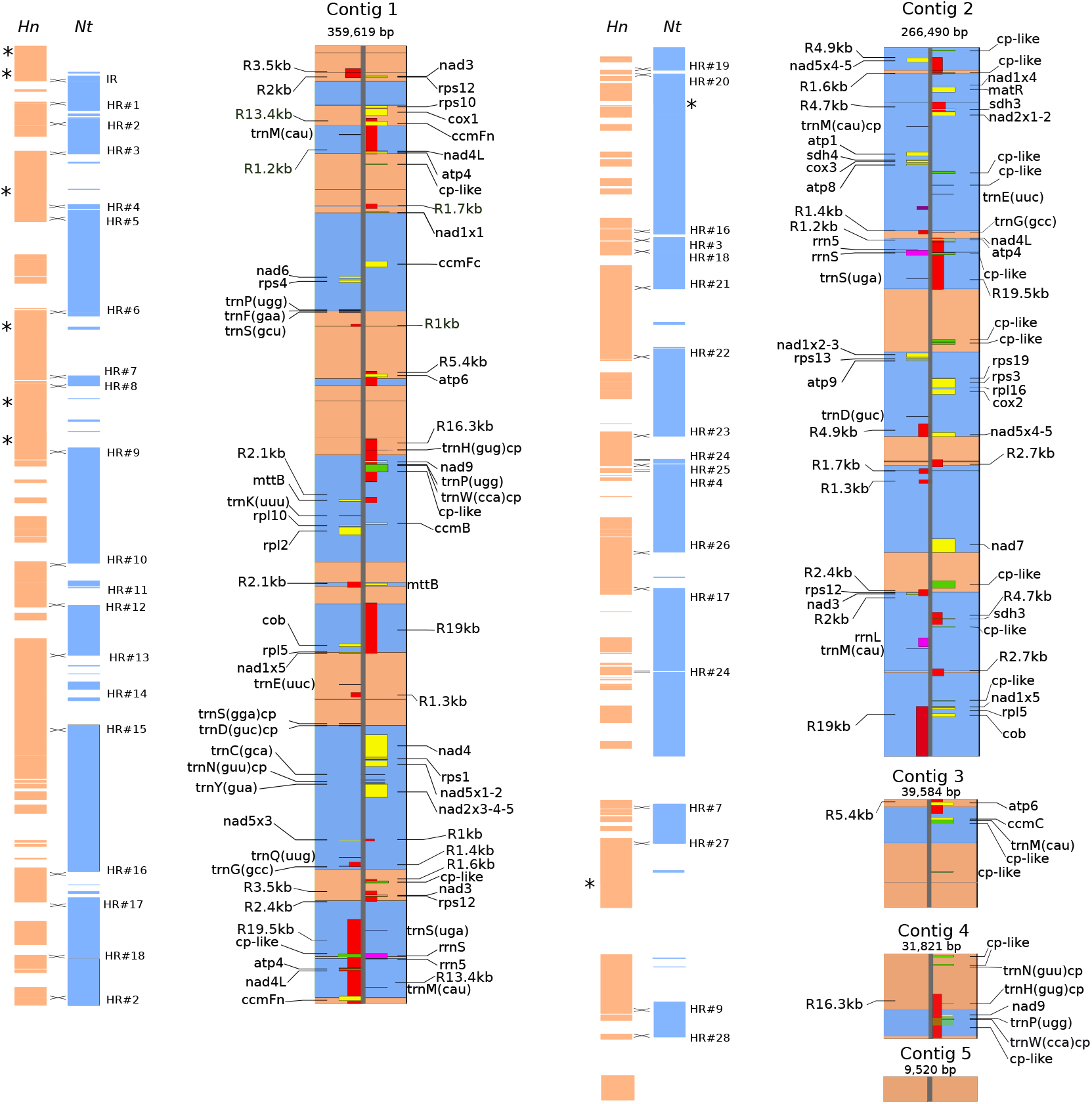
Mitochondrial genome of the repeated cybrid Nt[+Nt(+Hn)]. Scale maps of the five mitochondrial contigs totaling 707,034 bp. The alternate wide orange and blue boxes on the contig maps represent regions inferred to derive from the parents *H. niger* or *N. tabacum*, respectively. The cybrid contig maps show full-length genes, repeats >1 kb in length (red boxes labeled “R”, followed by the repeat length in kb), and chloroplast-derived regions > 300 bp in length (green boxes). Genes marked on different sides of the vertical lines are transcribed from opposite strands of the genome. The thinner orange and blue boxes to the left of the cybrid contig maps indicate regions of the cybrid mtDNA shared (as defined by BLAST hits) with the mitochondrial genomes of *H. niger (Hn)* or *N. tabacum (Nt)*, respectively. Crossing lines between BLAST hits indicate the 28 homologous (HR) and one illegitimate (IR) intergenomic recombination events, respectively. Intragenomic homologous recombination events are indicated with asterisks.

### Recombination events in the cybrid mitochondria

Once the parental mitochondrial genomes were united inside the cybrid mitochondria following mitochondrial fusion, they were able to undergo homologous recombination across the >1,000 regions of 100-9,000 bp with high identity between the two genomes. These homologous regions formed non-identical repeats (95-100% identity) in the initial cybrid mitochondria.

By comparing the cybrid and parental mtDNAs, we first identified the regions inherited from each parent based on sequence identity. We found 47 interspersed tracts of tobacco and *H. niger* with 99%-100% identity to the parental sequences (Figure 2). The analyses of the cybrid BLASTn hits to each parental mtDNA (orange and blue thin boxes on the left of each cybrid contig in Figure 2) indicated that the recombination events took place at homologous regions (with one exception) of variable lengths between the parental mtDNAs. We inferred 28 intergenomic homologous recombination events (HR#) giving rise to the chimeric Mv-1-1g mtDNA (Figure 2). The estimated length of the parental recombining regions was 2,500 bp in average, ranging from 205 to 9,850 bp and 94-100% sequence identity (Table 1). Based on the estimated minimal length of homology required by the enzyme RecA (Hua et al., 1997; Shen and Huang, 1986; Sugawara et al., 2000), one case of illegitimate recombination was inferred, in which the parental recombining regions shared only 19 bp (Figure 2 and Table 1).

**Table 1:** “Events of recombination between the parental mitochondrial genomes *(Nicotiana tabacum* and *Hyoscyamus niger)* in the cybrid Mv-1-1g.

| Location of recombination |  |  | Estimated mechanism of recombination |
| --- | --- | --- | --- |
| Contig coordinates | within a repeat? | within a coding region? |  |
| Contig 1: 13264-13280 |  | no | Illegitimate recombination through MMBIR |
| Contig 1: 22281-22322 |  | no | BIR or dHJ resolved through CO |
| Contig 1: 29413-29969 and 356835-357391 | Repeat 13.4kb | ccmFn | BIR or dHJ resolved through CO |
| Contig 1: 40094-40414 and Contig 2: 71414-71858 | Repeat 1.2kb | atp4; nad4L | dHJ resolved through CO |
| Contig 1: 59848-60456 and Contig 2: 159047-159516 | Repeat 1.7kb | no | dHJ resolved through NCO |
| Contig 1: 62209-62755 |  | nad1x1 | BIR or dHJ resolved through CO |
| Contig 1: 99430-99589 |  | trnF | BIR or dHJ resolved through CO |
| Contig1: 124863-125009 and Contig 3: 2760-2906 | Repeat 5.4kb | no | BIR or dHJ resolved through CO |
| Contig 1: 127505-127569 | Repeat 5.4kb | no | BIR |
| Contig1: 153523-153668 and Contig 4: 20699-20844 | Repeat 16.3kb | no | BIR or dHJ resolved through CO |
| Contig1: 193785-194096 |  | no | BIR or dHJ resolved through CO |
| Contig1: 201402-202870 | Repeat 2.1kb | mttB | SDSA or dHJ dissolution. |
| Contig1: 209565-209681 | Repeat 19kb | no | BIR |
| Contig 1: 227466-227797 | Repeat 19kb | rpl5 | BIR |
| Contig1: 244829-245456 | Repeat 1.3kb | no | SDSA or dHJ dissolution |
| Contig 1: 255104-255156 |  | no | BIR or dHJ resolved through CO |
| Contig1 :309198-309283 and Contig 2: 68948-69033 | Repeat 1.4kb | no | dHJ resolved through CO |
| Contig1: 320931-320984 and Contig 2: 204640-204666 | Repeat 2.4kb | no | dHJ resolved through CO |
| Contig1: 341742-342696 and Contig 2: 76106-77060 | Repeat 19.5kb | rrnS | SDSA, dHJ resolved through NCO, or dHJ dissolution |
| Contig 2: 8543-8624 | Repeat 1.6kb | no | BIR |
| Contig 2: 9992-10107 | Repeat 1.6kb | no | BIR |
| Contig 2: 90553-90835 | Repeat 19.5kb | no | BIR |
| Contig 2: 113811-114537 |  | no | BIR or dHJ resolved through CO |
| Contig 2: 145963-146237 | Repeat 4.9kb | nad5x5 | BIR |
| Contig 2: 155449-155768 and 234253-235034 | Repeat 2.7kb | no | dHJ resolved through NCO |
| Contig 2: 156753-157001 | Repeat 2.7kb | no | BIR |
| Contig 2: 189075-189866 |  | nad7x4 | BIR or dHJ resolved through CO |
| Contig 3: 15707-16024 |  | no | BIR or dHJ resolved through CO |
| Contig 4: 30891-30974 | Repeat 16.3kb | no | BIR |

Overall, the recombination events were unevenly distributed across the cybrid mtDNA (Figure 2). The two farthest consecutive points of recombination were located at ~54 kb from each other (HR#15 and HR#16), and the closest ones were positioned at a distance of 1,268 bp (Figure 2 and Table 1). Of the 29 intergenomic recombination events, eleven took place within or near coding regions resulting in five chimeric genes (see below).

In addition, we identified eight intragenomic homologous recombination events in the cybrid mtDNA, *i.e*. recombination between two regions derived from the same parent (asterisks in Figure 2). The recombining regions are repeats of estimated average length 510 bp, ranging from 73 to 2,623 bp and 86-100% identity. Noticeably, most of the intragenomic recombination events (87.5%) took place within sequences of the *H. niger* mtDNA. This might be explained by the high frequency of repeats in the *H. niger* mtDNA that could generate alternative genomic configurations through recombination events (Sanchez-Puerta et al., 2015).

### Parental contribution to the cybrid mitochondrial genome

Overall, 63% and 37% of the cybrid mtDNA were inherited from tobacco and *H. niger*, respectively. To examine the parental composition of the cybrid mtDNA in more detail, we distinguished three classes of sequences that contributed to the formation of the cybrid mitochondria: i) those shared by the two parents, *i.e*. homologous regions that included all known genes and many intergenic sequences; these sequences turned into non-identical repeats when the parental mtDNAs recombined to form the cybrid; ii) those exclusive to *H. niger*, and iii) those exclusive to *N. tabacum*. The repeated cybrid genome conserved one copy of almost all the *N. tabacum* exclusive sequences (99%) and 82% of those exclusive of *H. niger*. Overall, only 56 kb (11%) of the non-repeated sequences were eliminated. In stark contrast, 194 kb (~48%) of the repeated sequences were lost (Figure 3), leaving at least one copy of the homologous sequences in the cybrid Mv-1-1g. Of the 203 kb non-identical repeats initially present in the cybrid mitochondria (totaling 406 kb), only 10.5 kb sequences were retained from both parents (including 3 genes), and 172 kb and 19.1 kb were maintained from *N. tabacum* or *H. niger*, respectively (Figure 3). Most coding sequences (yellow areas in Figure 3) were retained from *N. tabacum*.

**Figure 3.**
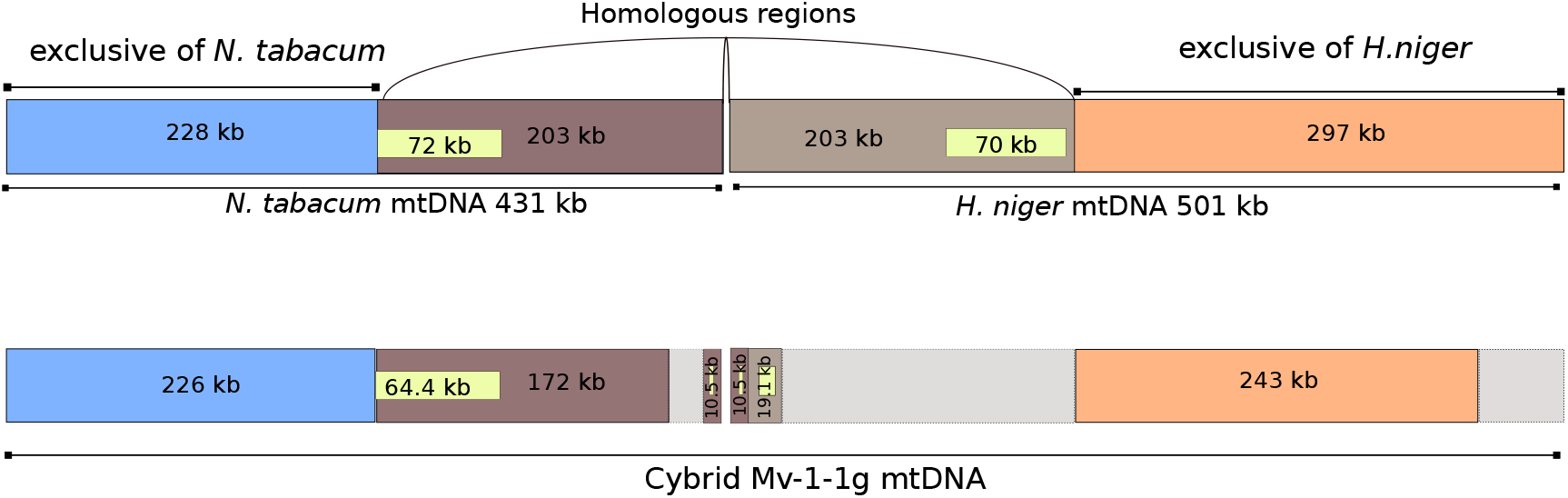
Origin of the cybrid mtDNA with respect to its parental composition. Top: schematic representation of the parental mitochondrial genomes (*Nicotiana tabacum* and *Hyoscyamus niger*) that contributed to the formation of mtDNA in the repeated cybrid Nt [+Nt(+Hn)] through two protoplast fusions. Total amount of exclusive and shared parental sequences are drawn to scale. Bottom: summary status of the composition of the cybrid mtDNA. Colored regions indicate sequences derived from tobacco- (blue and dark brown) or *H. niger*- (orange and light brown). Yellow regions represent coding sequences. Parental regions not present in the cybrid are shown in grey. Novel repeats in the cybrid mtDNA are not included, which explains the difference between the total length of the cybrid and the estimated amount of homologous and exclusive sequences derived from each parental mitochondria.

If we consider the double contribution of the tobacco mitochondria in the two successive protoplast fusions, the overall loss of repeated sequences increases significantly. The whole length of the tobacco mtDNA should be considered as a repeat and because the tobacco-derived regions were not found twice in the cybrid mtDNA (with very few exceptions), then, an extra 431 kb of repeated regions were also lost along the whole process.

### Mitochondrial gene content

Without counting repeats, the mtDNA of Mv-1-1g contained genes for 37 proteins, three rRNAs, and 21 tRNAs, i.e. the same number of mitochondrial genes as in the parental lines. The tRNA gene content is identical to tobacco, and not to *H. niger*, because the gene *trnD* is present while the gene *trnL-cp* is absent. In addition, the Mv-1-1g genome had two full-length copies of 11 genes (including the gene regions nad1_exon5 and nad5_exons4 and 5), and four genes were triplicates. Seven of these multi-copy genes resulted from the retention of full-length copies of each parent (*nad1×5, nad3, rps12*) or through the formation of chimeric genes originated from both parents (*atp4, ccmFn, rp15, rrnS*). The other duplicates were inherited from a single parent: *atp6, cob, nadθ, mttB, rrn5*, and *sdh3* (Table 2).

**Table 2:**
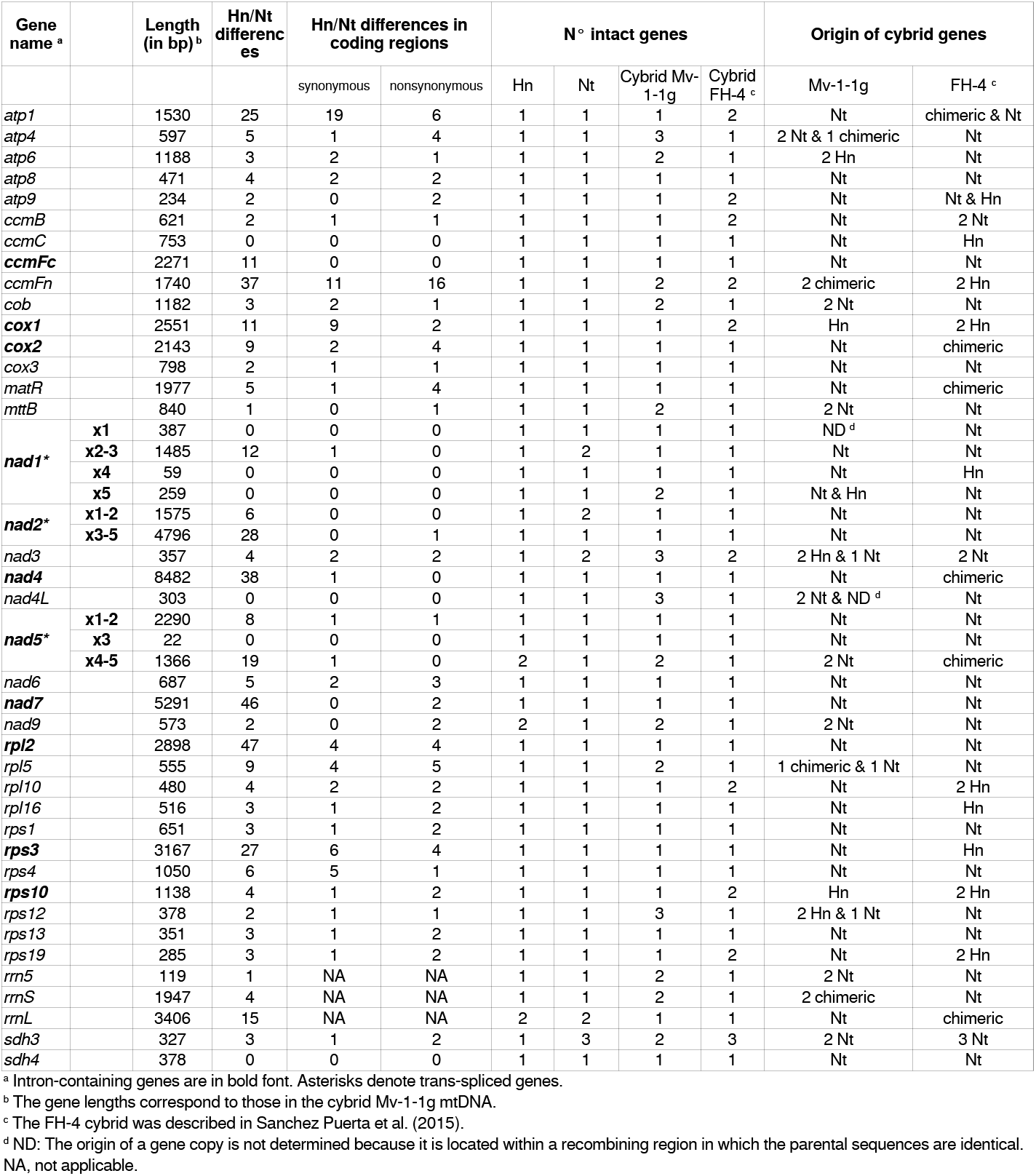
“Comparison of gene content in the mitochondrial genomes of the parents *(Nicotiana tabacum* (Nt) and *Hyoscyamus niger* (Hn)) and in the cybrids between Nt and Hn (Mv-1-1g and FH-4).

The protein-coding chimeric genes in the mtDNA of Mv-1-1g, *ccmFn* (two identical copies), *atp4*, and *rp15*, had nonsynonymous differences in respect of the parents, so the translated proteins would also be chimeric (Figure 4a). The genes *ccmFn* and *atp4* produced chimeric proteins that had eight or two different amino acids compared to the proteins in either parent, respectively. Finally, *rp15* showed two or three nonsynonymous changes compared to *H. niger* or tobacco, respectively (Figure 4a). Two of the genes that were inherited from both parents (*nad3, rps12*) had nonsynonymous differences between them and similar hydrophobicity profiles (Figure 4b), while *nad1×5* was identical in both parents.

**Figure 4.**
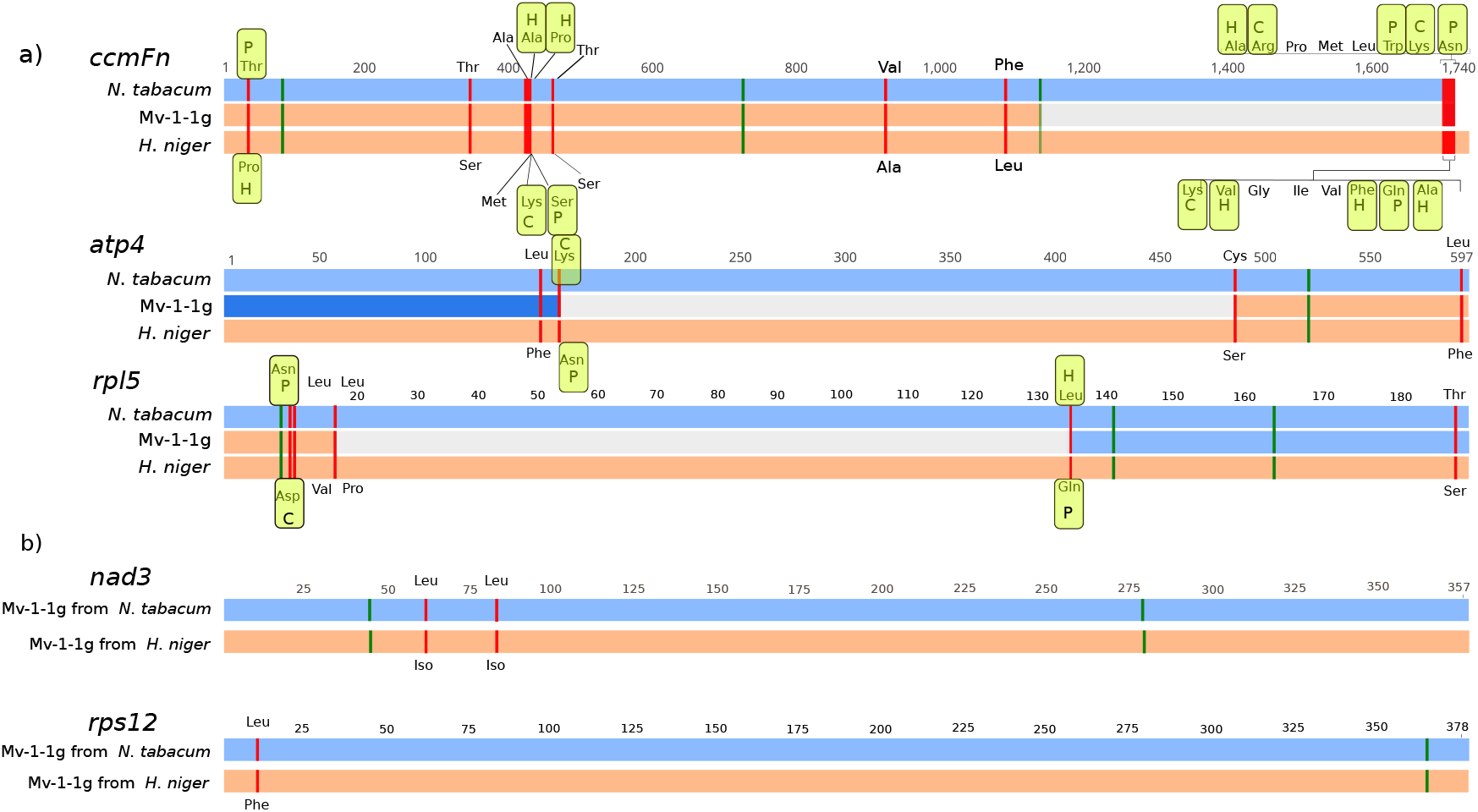
Schematic representation of differences in chimeric and two parent-derived genes in the cybrid Mv-1-1g mtDNA with respect to the parental sequences. a) Chimeric genes (*ccmFn, atp4*, and *rp15*) in the cybrid Mv-1-1g formed by intergenomic recombination of parental mitochondria. b) Genes in the cybrid (*nad3, rps2*) found in two copies, one derived from each parent. Blue and orange horizontal boxes indicate sequences derived (or from) *N. tabacum* and *H. niger*, respectively. Gray boxes depict crossover regions in which the parental sequences are identical. Red and green marks indicate nonsynonymous and synonymous differences, respectively, between the parental lines. The encoded amino acid is shown for the nonsynonymous differences. Amino acids with hydrophobic (H), polar (P), or charged (C) side chains are indicated within yellow rectangles where the nonsynonymous differences implicate a change in the amino acid chemical group. Numbers denote sequence length in bp.

Of those inherited from a single parent, 27 protein-, two rRNA-, and 14 tRNA-genes originated exclusively from *N. tabacum*, while only three protein and two tRNA genes derived exclusively from *H. niger* (Table 2). The three protein genes inherited only from *H. niger* (*atp6, cox1*, and *rps10*) had nonsynonymous differences from those in *N. tabacum* and different hydrophobicity patterns. In addition, the gene *cox1* had an intron in *H. niger* and not in tobacco. Also the aforementioned genes had differences in the predicted editing sites but these differences are functionally irrelevant because *H. niger* genes had a subset of the tobacco editing sites.

### ORFs in the cybrid mtDNA

We searched for uncharacterized ORFs in the cybrid mtDNA that could be related to the CMS condition and the atypical flower phenotype. In the cybrid mtDNA those ORFs inherited exclusively from *H. niger* are now in a foreign nuclear background, and could be involved in the CMS phenotype (Leino et al., 2005). We identified 66 ORFs derived from *H. niger* that did not show sequence similarity to ORFs in *N. tabacum* mtDNA. Two of those ORFs showed one of the most common features found in CMS-related genes, i.e. they start with partial sequences of known mitochondrial genes followed by an uncharacterized region (Chen and Liu, 2014). One of them, ORF281 (located in contig 1) of 843 bp, begins with the first 66 bp of the gene *rps13*. The other, ORF289 (located in contig 2) of 867 bp, begins with the first 173 bp of the gene *nad9*.

In addition, new ORFs could arise in the cybrid mtDNA by intergenomic recombination. These newly formed ORFs are considered to be good candidates for determination of CMS (Chase, 2007; Touzet and Meyer, 2014). We found two ORFs located at the sites of recombination of the cybrid Mv-1-1g mtDNA. ORF326 of 978 bp was a new ORF with a domain from DNA polymerase B2 superfamily identified by HMMER (Finn et al., 2015). ORF117 (located in contig 1) was chimeric and showed nonsynonymous substitutions to similar ORFs in the parental mtDNAs.

## Discussion

Somatic hybridization is a useful tool for bypassing the incompatibility barriers in sexual crosses of distinct plant species, as well as the barriers imposed by the uniparental mode of transmission of mitochondrial genes in sexual crosses of plants (Greiner and Bock, 2013). Protoplast fusion leads to *de novo* nuclear and cytoplasmic genome combinations, and plant hybrids almost invariably contain recombinant mitochondrial genomes (Akagi et al., 1995; Aleza et al., 2016; Belliard et al., 1979; Xiang et al., 2004). This turns somatic hybrids into an interesting subject for further studies of mitochondrial genome recombination, evolutionary dynamics, and nuclear-cytoplasmic coordination. Cybrids obtained via somatic hybridization might represent a valuable technological tool for transferring organelle-encoded traits to cultivars, such as CMS (Fitter et al., 2005; Sun et al., 2005). The CMS condition can result from the formation of chimeric mitochondrial genes, which are frequently formed in somatic hybrids (Chase, 2007; Kim and Kim, 2006). Little is known about the overall extent of mitochondrial recombination in somatic hybrids, the mechanisms involved, and the constraints operating on mtDNA recombination in the context of nuclear-cytoplasmic interactions and incompatibilities. The present study has been conducted in an attempt to advance our understanding of plant mitochondrial genome recombination, and their potential technological applications in plant breeding.

### Evaluating the genetic basis of the CMS phenotype

The origin and mechanisms of CMS are still mysterious (Charlesworth, 2017; Chase and Gabay-Laughnan, 2004; Hanson and Bentolila, 2004). Unusual ORFs determining CMS have been discovered in a wide range of plant species, showing considerable diversity of the proteins involved and the specific phenotypic effects (Chase, 2007). Most of the mitochondrial-encoded loci responsible for CMS that were reported to date shared two features: they are ORFs formed by sequences of unknown origin combined with pieces of standard mitochondrial genes and they are expressed because they are fused directly to a mitochondrial promoter region or they are cotranscribed with upstream mitochondrial genes (Carlsson et al., 2008; Chase, 2007; Chase and Gabay-Laughnan, 2004; Hanson and Bentolila, 2004; Schnable and Wise, 1998) In other cases, CMS genes were not chimeric and, instead, contained sequences from a single source (Bonhomme et al., 1992; Ducos et al., 2001; Okazaki et al., 2013).

The cybrid analyzed in this work manifested CMS, and this condition was maintained after a number of backcrosses with *N. tabacum* as well as after a second fusion event with a different nuclear background of *N. tabacum*. These experiments eliminated nuclear-factors as the cause of CMS (Zubko et al., 1996). Instead, the origin of such important feature should be associated with changes in the mitochondrial genome, or resulted from nuclear-mitochondrial incompatibilities. In this work, we found unusual ORFs and features in the mitochondrial genome that could be responsible for the CMS phenotype in the Mv-1-1g repeated cybrid. For example, the mtDNA of the cybrid Mv-1-1g contained respiratory complexes I, IV, and V formed by subunits encoded by different combinations of *H. niger*-derived (*i.e. nad1, nad3, atp6, cox1*) and/or chimeric (i.e. *atp4*) mitochondrial genes, together with tobacco-derived genes (*i.e. nad2, nad4, nad5, cox2, cox3, atp1, atp8, atp9*) in a tobacco nuclear background. The assembly of protein complexes formed by genes of different origin, carrying new mutations, could cause steric incompatibility, reduction of substrate affinity, and prevention of activity of respiratory complexes, causing CMS (Ducos et al., 2001; Pineau et al., 2005; Rand et al., 2004).

Despite containing CMS genes, plants could be male fertile due to the presence of restorer genes in their nuclear genomes. When these pre-existing CMS genes are moved to a novel nuclear background, through genetic crossing or somatic hybridization, suppressors are removed and CMS is expressed (Carlsson et al., 2008; Leino et al., 2005). In the mtDNA of the Mv-1-1g cybrid, there are 66 ORFs derived from *H. niger* that do not have similarity with ORFs in *N. tabacum* mtDNA. Even though *H. niger* is male fertile, these ORFs could be involved in the CMS phenotype observed in the repeated cybrid. Of the 66 ORFs derived from *H. niger*, two were chimeric, beginning with a short piece of the genes *rps13* and *nad9*, respectively, and represent worthy candidates for CMS. Variety of new ORFs and other reconstructed mtDNA regions implies a unique avenue for future work towards clarification of their roles in particular phenotypic peculiarities of CMS flower composition and other developmental features of cybrids. Extended investigation of links between mtDNA changes and various phenotypic traits in cybrids derived from the same remote parents might bring in depth understanding of nucleo-cytoplasmic communications as well as a potentially new source of genetic variation for evolution and breeding.

### Recombination events in the cybrid mitochondria

Plant mtDNAs undergo frequent recombination processes that have central roles in DNA repair and replication (Backert and Borner, 2000; Kajander et al., 2001; Ling et al., 1995). Studies on plant mitochondrial recombination and repair have focused mainly on *Arabidopsis* nuclear mutants that described the role of nuclear-encoded genes, such as OSBs (Zaegel et al., 2006) and homologs of the bacterial genes *MutS* and *recA* (Arrieta-Montiel et al., 2009; Davila et al., 2011; Miller-Messmer et al., 2012; Shedge et al., 2010). Rearrangements in plant mtDNAs can be explained by the action of homologous recombination (HR) pathways or through illegitimate recombination involving sequence microhomologies (Kubo and Newton, 2008; Kühn and Gualberto, 2012). Several studies suggested that the minimal length of the homology region required for HR is 24-40 bp depending on the organism although its efficiency increases rapidly with the size of the homologous regions that undergo recombination (Hua et al., 1997; Shen and Huang, 1986; Sugawara et al., 2000).

Double strand breaks (DSBs) arise from exogenous and endogenous sources and can be repaired by three major HR mechanisms (Mehta and Haber, 2014). A DSB leads to 5’->3’ degradation of the broken ends resulting in 3’ single-stranded DNA tails (Figure 5). The activation of different pathways to repair the DSB depends on the structure of the broken ends of the DNA and the factors available (Mehta and Haber, 2014). In the double Holliday Junction (dHJ, also known as Double Strand Break Repair DSBR) pathway, two Holliday junctions are formed and, depending on the resolution, it can result in crossover (CO) or non-crossover (NCO) products (Resnick, 1976; Szostak et al., 1983). Alternatively, the dHJ can be dissolved via producing gene conversion products (Wu and Hickson, 2003). In the Single-Dependent Strand Annealing (SDSA) pathway, strand invasion is rejected and the 3’ ssDNA tail aligns with the other end of the DSB that results in a faithful repair without crossover products, leading to gene conversion (Paques and Haber, 1999). Break Induced Replication (BIR) plays an important role in the repair of the one-ended DSBs that can be formed as result of a collapse or arrest of the replication fork or when only one end of the DSB shares homology with a donor sequence (Anand et al., 2013). Strand invasion occurs and the synthesis proceeds to the end of the DNA molecule, resulting in a non-reciprocal crossover (Anand et al., 2013; Malkova et al., 1996, 2005).

**Figure 5.**
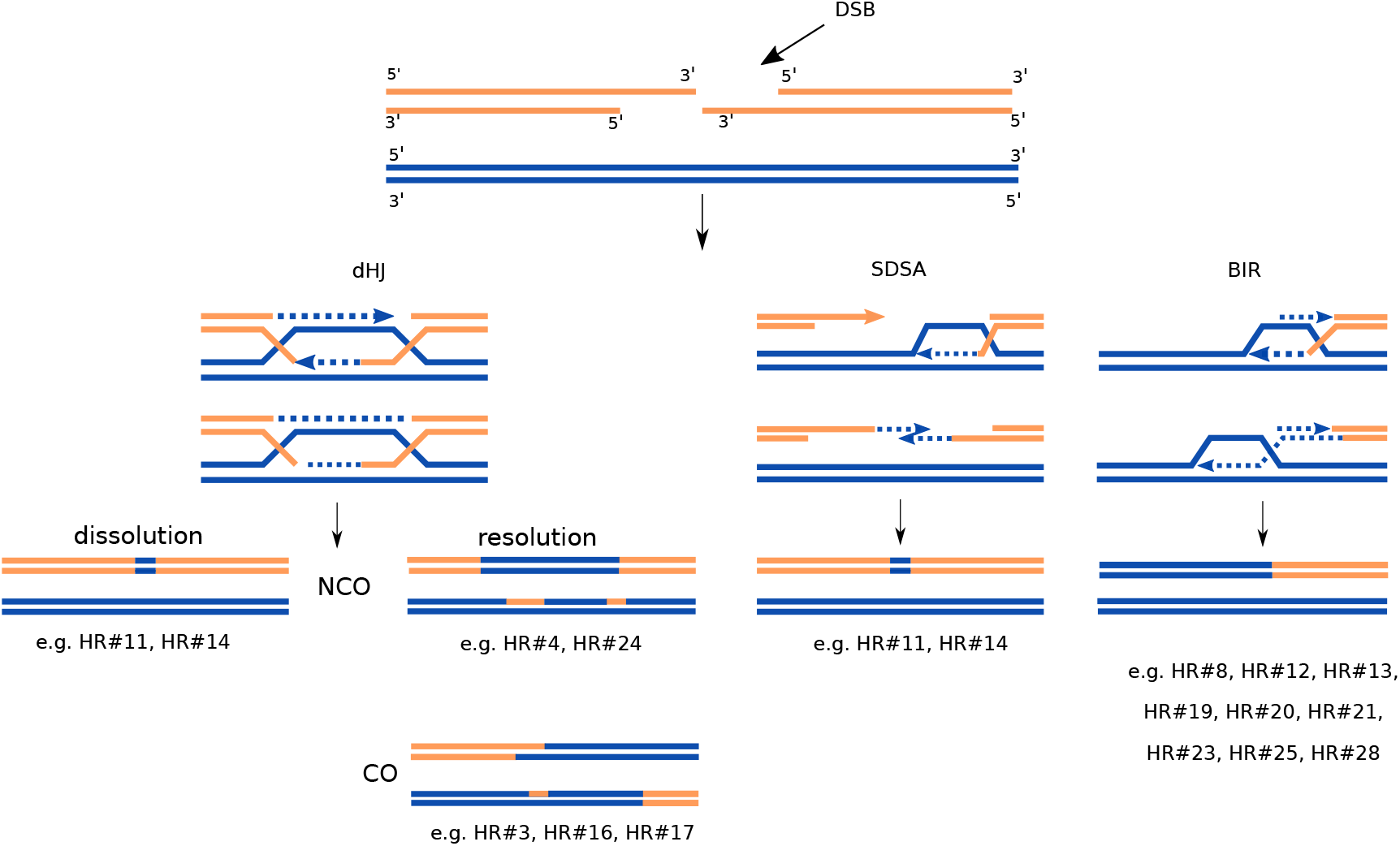
Homologous recombination pathways. Mechanisms of DSB repair. DSBs are 5’ digested resulting in 3’ single-stranded DNA tails. The double Holliday Junction (dHJ) pathway involves two strand invasions and can be resolved either by helicase and topoisomerase-mediated dissolution resulting in noncrossover (NCO) products or cleaved by HJ resolvases to produce both crossover (CO) or NCO outcomes. In the Synthesis Dependent Strand Annealing (SDSA) pathway, the newly synthesized strand dissociates from the D-loop and results in a NCO outcome with no change to the template DNA. The Break Induced Replication (BIR) pathway involves both leading and lagging strand synthesis and results in the loss of heterozygosity or, a nonreciprocal translocation (if the template is located ectopically). Newly synthesized DNA is depicted as dashed lines in the same color as the template; arrowheads indicate direction of DNA synthesis. Examples of inferred homologous recombination pathways in the cybrid Mv-1-1g mtDNA are indicated. Blue and orange horizontal lines indicate sequences derived or from *N. tabacum* and *H. niger*, respectively.

At the time of protoplast fusion, the set of homologous sequences (> 1,000 regions) between the two parental mtDNAs became non-identical repeats allowing the process of intergenomic HR. We analyzed the products of recombination between the two parental genomes to infer which mechanisms took place during cybrid formation (Table 1, Figure 5). We identified three events of symmetric recombination produced by the dHJ pathway resolved towards CO (HR#3, HR#16 and HR#17), nine cases of non-reciprocal recombination (HR#8, HR#12, HR#13, HR#19, HR#20, HR#21, HR#23, HR#25, HR#28) indicating the involvement of the BIR pathway, and two events that could be explained by dHJ with NCO resolution (HR#4 and HR#24). Two cases (HR#11 and HR#14) could be explained either by SDSA or by dHJ dissolution. In addition, 11 single products of recombination were present in the cybrid mtDNA, for which the recombination pathway could not be determined (Table 1).

Overall, BIR was the most frequent mechanism in agreement with previous observations in another cybrid (Sanchez-Puerta et al., 2015). This could be due to frequent occurrence of stalled replication forks (Anand et al., 2013). Alternatively, a DSB close to the end of a homologous region in the parental genomes would lead to BIR because one end will not find a homology region to repair the DSB by gene conversion. BIR is a highly mutagenic pathway because it can result in chromosomal rearrangements and the loss of a large piece of DNA (Sakofsky et al., 2012). It has been suggested that BIR is suppressed when a DSB has two ends and a more conservative HR pathway is followed (Llorente et al., 2008).

In addition to HR, there are mechanisms to repair DSB that require very short or no homology regions. Recombination at microhomologies causes illegitimate recombination through three molecular pathways: Non Homologous End Joining, (NHEJ), Microhomology-Mediated End Joining (MMEJ), and Microhomology-Mediated Break Induced Replication (MMBIR) (Hanson and Bentolila, 2004; Hastings et al., 2009; Kubo and Newton, 2008; Schnable and Wise, 1998). NHEJ is the predominant repair mechanism for DNA DSB in the nuclear genome (Chang et al., 2017). This mechanism directly ligates broken DNA strands, without 5’ resection, and requires microhomologies of 1-4 bp (Lieber, 2010). This pathway can produce blunt joints or small deletions or insertions in the breakpoint junction. MMEJ starts with the digestion of 5’ DNA strand to obtain 3’ DNA tails. The exposed microhomologies anneal and the gaps are filled. This mechanism results in deletion of the DNA regions flanking the original break; it is an error prone repair mechanism (Bennardo et al., 2008). MMBIR is a replicative microhomology mediated mechanism. In this model, the replication fork stalls with the following resection of 5’ ends, generating 3’ strand overhangs, which can invade a region with microhomology on a different DNA template to establish a new replication fork (Hastings et al., 2009)(Hastings et al., 2009). In this study, a single recombination across a short repeat was found with no deletions suggesting that it is the result of MMBIR.

### Parental content of the cybrid mtDNA

Several factors could shape the gene content, length, and parental origin of the mitochondrial genome of a cybrid plant. For example, the interactions of the recombinant mitochondria with the chloroplast (derived from *H. niger*, in this case) and the nuclear (derived from tobacco) genomes could impose constraints on the number and origin of the mitochondrial genes. Furthermore, the recombination pathways that may take place between the parental mtDNAs would influence the structure, chimerism, repeat content, and length of the cybrid mitochondrial genome. In a previous study, the mtDNA of a cybrid between tobacco and *H. niger, Nt(+Hn)* FH-4, had preferentially eliminated repeated sequences and retained a single allele (mostly from tobacco) for each mitochondrial gene (Sanchez-Puerta et al., 2015) Recently, the mtDNA of a cybrid between *Brassica napus* and *Raphanus sativa* also showed similar features (Arimura et al., 2018).

In the cybrid under study, a total of three mitochondrial genomes (two of tobacco and one of *H. niger*) were joined through two protoplast fusion events. Instead of a potentially expected giant mtDNA, the resulting cybrid mitochondria contained a genome that was 1.4-1.6 times greater than that of either parent, in agreement with observations in the other two cybrids mentioned above (Arimura et al., 2018; Sanchez-Puerta et al., 2015). In all cases, the cybrid mtDNA lost a substantial fraction of the parental mitochondrial genomes but it was larger than either parental mtDNA. The three cybrids also shared a high similarity in the parental composition of the mtDNAs. They maintained preferentially mitochondrial sequences of the same parent as the nuclear genome.

In particular, we found that most coding sequences in the cybrid Mv-1-1g were inherited from *N. tabacum*, as expected, given that the nuclear genome derived from tobacco and that the cybrid received two doses of mtDNA from *N. tabacum*. However, the cybrid mtDNA retained six genes or gene regions from *H. niger*. Three of them were maintained as single copy genes *(atp6, cox1, rps10)* and the other three as multi-copy genes with one allele derived from each parent (*nad1×5, nad3, rps12*). Similarly, in the cybrid Nt(+Hn) FH-4 previously analyzed (Sanchez-Puerta et al., 2015), 10 genes or gene pieces from *H. niger* were retained, including the genes *cox1* and *rps10*.

The presence of the chloroplast genome from *H. niger* in both cybrids could influence the retention of those genes. It is possible that the transferred chloroplasts may need genetically related mitochondrial sequences to be functional in a foreign nuclear background. Several reports described the close relationship of the organellar genomes and the cross-talk between organelles and the nucleus to maintain a tight cellular organization. Chloroplast and mitochondria show a high level of metabolic interdependency and a coordinated work, and this communication has been established during evolution (Leister, 2005; Raghavendra and Padmasree, 2003; Sabar et al., 2000; Smith and Keeling, 2015). Alternatively, the retention of a few genes from *H. niger* could be the result of genetic drift.

Besides the type of parental allele retained by each cybrid mtDNA, the three cybrids maintained a single allele of each gene, either from one parent or from the other, but rarely from both. According to Sanchez Puerta *et al*. (2015), this observation could be explained by a neutral mechanism that eliminates preferentially repeated sequences, or by adaptive forces given the putative deleterious effect of keeping potentially conflicting copies of coding regions derived from both parents.

A comparison of homologous and non-homologous regions inherited by the cybrids between tobacco and *H. niger* (Mv-1-1g and FH-4) showed that almost all regions that were exclusive of each parent were maintained (~89%), in stark contrast to the limited retention of those homologous sequences contributed by both parents (~48%) (Figures 5 and 6). This is also true for the cybrid between *B. napus* and *R. sativus* (Arimura et al., 2018) that retained 100% and 45% of the exclusive mitochondrial sequences from *B. napus* and *R. sativa*, respectively, while homologous sequences were kept only from *B. napus* (except for 4 kb from *R. sativus* representing 2% of the shared sequences). We hypothesize that the non-homologous parental sequences were not eliminated in the cybrid mtDNA because they could not undergo intergenomic HR. Instead, DSBs in these regions were necessarily repaired using another copy of the same parental genome. These non-homologous regions were non-coding DNA (except for uncharacterized ORFs) and would be evolving mainly by genetic drift.

**Figure 6.**
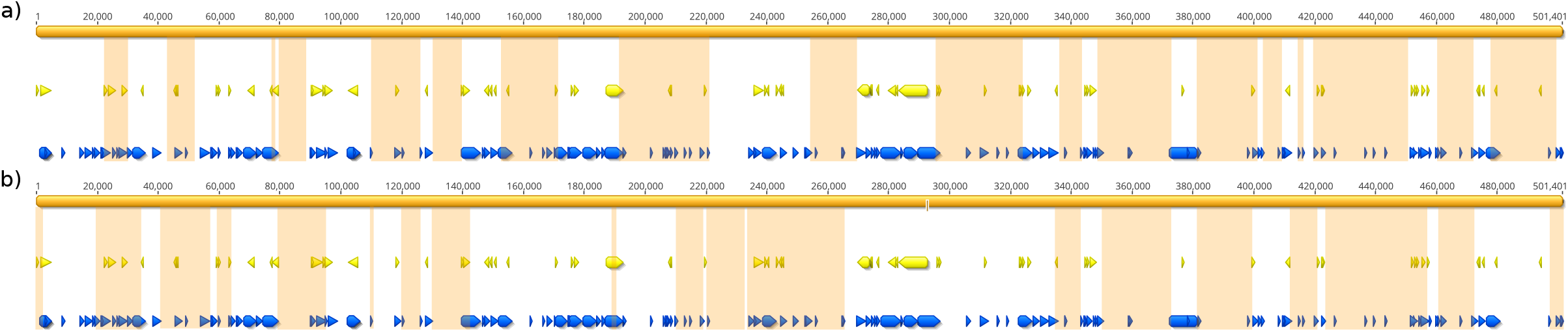
Regions of the parent *Hyoscyamus niger* retained in the cybrids. A linear map of the *H. niger* mitochondrial genome (in orange) shows the regions (in light orange) that were retained in the cybrids Mv-1-1g (a) and FH-4 (b). Yellow and blue arrows indicate coding sequences and shared regions between tobacco and *H. niger* mitochondrial genomes, respectively.

Of the homologous sequences in the parental mtDNAs, the cybrid Mv-1-1g maintained only a small fraction from both parents, while the majority was retained from a single parent (mostly tobacco). The homologous sequences include coding (34%) and non-coding (66%) sequences. Under the neutral model of evolution, we expected to find an almost unbiased parental composition of the non-coding homologous regions (derived randomly from *H. niger* o *N. tabacum)* in the cybrid mitochondria. Instead, we observed that, in all three cybrids mentioned above, the vast majority of the non-coding homologous sequences derived from the same parent as the nuclear genome (tobacco or *B. napus*, depending on the cybrid). This bias raises questions regarding the importance of non-coding sequences for the mitochondrial function/homeostasis/gene expression/nuclear cytoplasmic compatibility. Alternatively, the adaptive elimination of coding regions from *H. niger* could have dragged the surrounding non-coding regions along. In fact, most of the non-coding homologous sequences are interspersed among coding regions (Figure 6). Therefore, the putatively advantageous loss of *H. niger* genes from the cybrid mtDNA described above could be responsible for the deletion of the homologous non-coding regions of *H. niger* surrounding those genes.

Several recombination pathways could cause the elimination of genomic sequences, such as SSA (Sugawara et al., 2000; de Zamaroczy et al., 1983), recombination across short repeats, recombination via sub-stoichiometric intermediates (Small et al., 1989), and BIR (Malkova et al., 1996). Single-strand annealing (SSA) repairs DNA breaks that are flanked by long direct DNA repeats leading to a loss of DNA between the repeats (Sugawara et al., 2000). We did not find evidence for this DSB repair pathway in the cybrid mtDNA. Recombination via sub-stoichiometric intermediates could also explain deletions in plant mitochondrial genomes (Small et al., 1989), but we could not test this hypothesis directly. We propose that BIR could be one of the mechanisms responsible for the loss of the majority of the repeated regions derived from *H. niger*. Of the nine recombination events produced by BIR, seven implied the invasion of the *H. niger* strand into a tobacco-derived DNA, resulting in the loss of a homologous segment derived from *H. niger* (Figure 5). In addition, half-crossover BIR, caused by aberrant processing of BIR intermediates, could result in the loss of one of the two chromosomes participating in the recombination events (Deem et al., 2008; Haber and Hearn, 1985). This could explain several of the single product recombination events.

We conclude that *H. niger* genes and the flanking homologous regions were eliminated from the cybrid through natural selection and that BIR was the molecular pathway responsible for the loss of such sequences.

## Acknowledgements

This work was supported by Universidad Nacional de Cuyo (Sectyp M033), Agencia Nacional de Promoción Científica y Tecnológica (grant number PICT1762) to M.V.S.P and by NSF (grant number 1062432) to Indiana University, which supports the computer cluster.

## Supplementary Information

**Figure S1. Total read depth of the mtDNA of the repeated cybrid Nt[+Nt(+Hn)].** Scale maps of the five mitochondrial contigs of the cybrid Mv-1-1g. Spikes of the read-depth result from mismapping of reads derived from the chloroplast genome onto large mitochondrial regions with high identity to chloroplast sequences.

