## Supplementary material for "Elucidating genomic patterns and recombination events in plant cybrid mitochondria": Suppl. Figure 1

Figure S1. Total read depth of the mtDNA of the repeated cybrid Nt[+Nt(+Hn)].
Scale maps of the five mitochondrial contigs of the cybrid Mv-1-1g. Spikes of the read-depth are the result of mismapping of reads derived from the chloroplast genome onto large mitochondrial regions with high identity to chloroplast sequences.

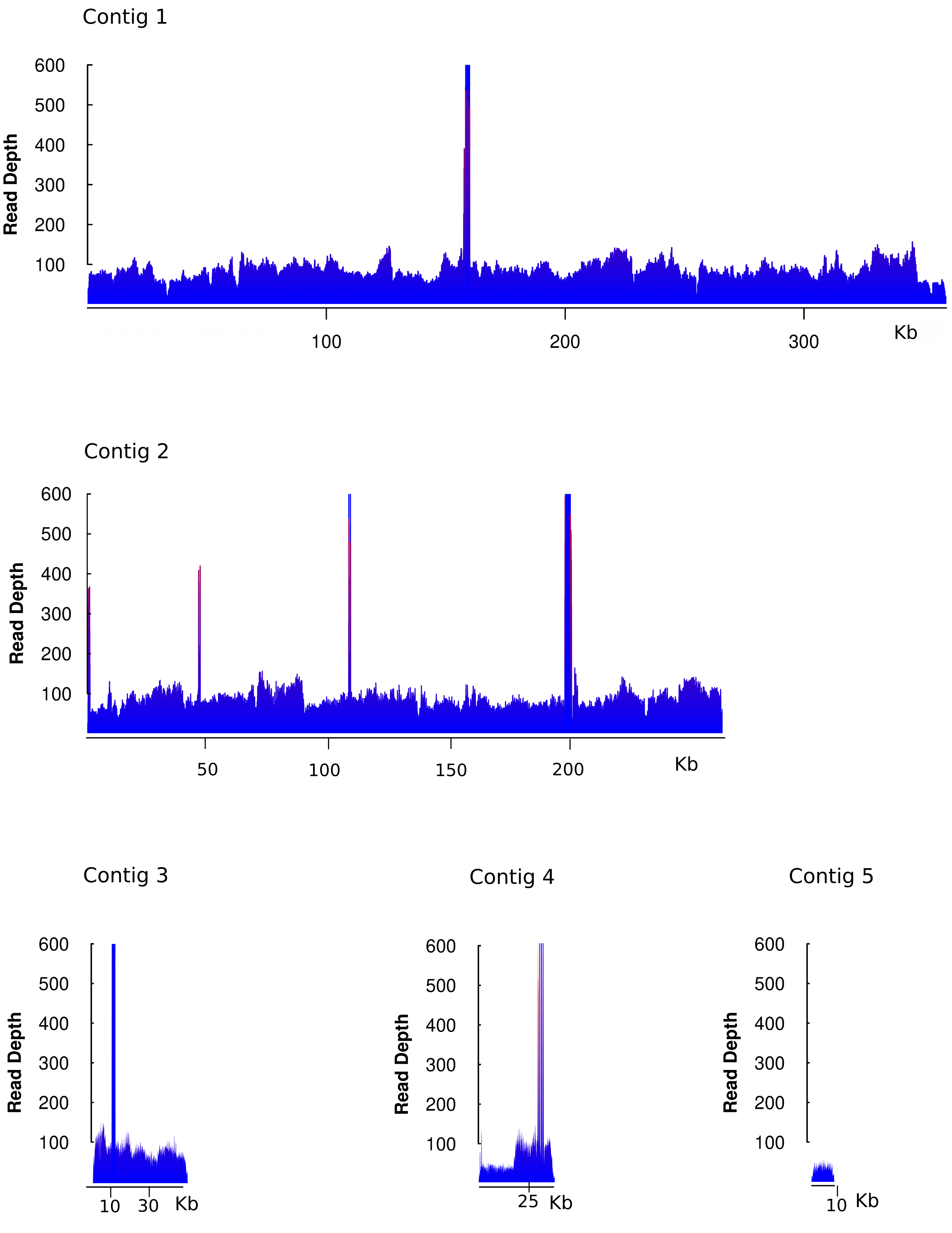
